# Rice OsML1, a distant plant homologue of animal MD-2 protein, can also bind to and recognize bacterial LPS and co-triggers innate immunity

**DOI:** 10.64898/2026.05.03.722507

**Authors:** Mengtian Pei, Xuze Xie, Stefan Olsson, Zonghua Wang, Wenwei Lin, Guodong Lu

**Affiliations:** State Key Laboratory of Ecological Pest Control for Fujian and Taiwan Crops, Fujian Agriculture and Forestry University, Fuzhou 350002, China; Haixia Institute of Science and Technology, Fujian Agriculture and Forestry University, Fuzhou 350002, China

**Keywords:** Oryza sativa, PITPs, OsML1, LPS receptor, MAMP, innate immunity

## Abstract

Lipopolysaccharides (LPSs) are pathogen-associated molecular patterns (PAMPs) of Gram-negative pathogenic bacteria recognized by plants, triggering typical pattern-triggered immunity (PTI) responses. However, a LPS sensing receptor for the recognition of plants remains largely undefined. A plant receptor for lipopolysaccharide (LPS) has not yet been identified. Here, we identify a plant protein, OsML1, with homologies to animal MD-2, which is capable of binding LPS. Furthermore, it may act as a molecular chaperone to assist CK1 in perceiving LPS signals. Our results show that OsML1 functions as an LPS-binding protein recognizing LPS and participates in downstream rice immune response activation. Structural modeling and sequence analysis revealed that OsML1 contains both a typical ML domain and a conserved three-dimensional β-barrel structure as mammalian MD-2 proteins. Microscale thermophoresis assays confirmed that OsML1 binds LPS with high affinity. Functional analyses further demonstrated that OsML1 knockout plants show reduced resistance to the rice bacterial blight pathogen, as well as attenuated ROS bursts upon LPS treatments, whereas overexpression plants show enhanced immune responses. Metabolomic profiling indicated significant metabolic changes in OsML1 knockout plants, particularly in immune-related pathways involving lipids, amino acids, and antimicrobial compounds. OsML1 is consequently a structurally conserved and functional LPS-binding protein linking lipid metabolism, LPS perception, immune activation, and metabolic regulation. Phylogenetic and structural analyses revealed that OsML1 likely arose from a duplication of OsML2, forming an independently functional subgroup within the PITP family. Our study identifies OsML1 as a LPS recognition factor involved in LPS sensing and downstream ROS bursts activation, callose deposition, and broad-spectrum gene expression of resistance. These findings expand our knowledge of bacterial LPS perception and immune regulation in plants, offering novel targets and strategies for disease-resistant breeding.

## Introduction

Rice is among the most significant staple foods, feeding nearly half of the world’s population. Despite significant efforts directed towards cultivating highly productive and multi-resistant rice varieties, the impact of diseases continues to cause substantial annual reductions in rice yield [1]. Due to its fixed growth environment and singular planting pattern, the rice plant is susceptible to various diseases and insects. To cope with this stress, rice has evolved complex and systematic resistance mechanisms.

Phosphatidylinositol transfer proteins (PITPs) are indispensable for the growth and development of eukaryotes [2]. The PITP Class IV ML has been extensively researched in both mammals and insects. The beta-rich-sheets predicted within their ML domains consist of multiple chains and are believed to facilitate various biological activities through their interactions with specific lipids [3]. MD-2 is the initial ML discovered in a human kidney cell line, serving as an extracellular binding chaperone for Toll-like receptor 4 (TLR4) [4]. The combination of MD-2 and TLR4 leads to enhanced recognition and binding of lipopolysaccharide (LPS), a component of bacterial cell wall, thereby inducing the expression of immune response-activating cytokines interleukin 10 (IL-10) and transforming growth factor-1 (TGF-1) [5–7]. Numerous investigations have indicated the involvement of ML protein in the innate immunity process. LPS-binding activity has been observed in MD-2 of *Mus musculus*, MsMl-1 of *Manduca sexta*, and LvMl of *Litopenaeus vannamei*, suggesting that ML recognizes LPS signals and takes part in the innate immunity process [8–10]. The AgMdl-1 homolog of Anopheles gambiae plays a role in the immune defense of mosquitoes against Plasmodium falciparum infection [11]. The ML domain-containing protein BmEsr16 from the silkworm (Bombyx mori) can bind to a range of bacterial cell wall components, including LPS, lipid A, peptidoglycan (PGN), and lipoteichoic acid (LTA). Loss of function of BmEsr16 can result in a reduction of antibacterial response [12,13]. Furthermore, the salivary glands and hemolymph of castor seed ticks (*Ixodes ricinus*) express the IrML gene throughout all developmental stages, indicating its potential involvement in innate immune responses [14].

In our previous paper [2], we identified OsM1 as a possible MD2 protein in rice. Since MD-2 is a key recognition factor in the animal innate immune system that recognizes LPS, the discovery marks a key step in understanding the molecular mechanism of animal recognition of Gram-negative bacteria. MD-2 interacts with TLR4 to form a functional complex, mediating high-affinity recognition of LPS, thereby initiating downstream inflammatory signaling pathways. It is a core component of TLR4-dependent immune activation. This mechanism not only provides an antibacterial immune response for mammals, but also provides a molecular basis, but also provides a classic model for revealing how hosts recognize pathogen-associated molecular patterns. However, unlike the in-depth analysis of MD-2 function in animal systems, the proteins recognizing lipopolysaccharides by plants have been lacking for a long time, although studies have shown that LPS can induce reactive oxygen species bursts and calcium in plants. These are typical innate immune responses combined with other responses such as ion flux changes, but if a functional equivalent to animal MD-2 is involved in LPS recognition in plants, such as rice is yet to be known [4].

The lipid recognition factor and its mechanism of action have not yet been clearly concluded. In our previous paper, we identified OsM1 as a possible MD2 protein [2]. In this paper, we use a range of analyses, including bioinformatics and various experimental techniques, and show that OsML1 is a protein with similar LPS-binding and function in rice immunity, similar to what MD-2 has in animal systems. It is thus the first LPS receptor identified in plants.

## Materials and methods

### Plant materials and generation of transgenic rice plants

The rice cultivar used in this study was the wild-type line ZH11. Rice plants were grown in a controlled-environment chamber under the following conditions unless otherwise specified: 26°C, 50% relative humidity, and an 18 h light/6 h dark photoperiod.

Competent cells of the yeast strain AH109, Escherichia coli DH5α, and E. coli BL21(DE3) were prepared and stored in our laboratory. Xanthomonas oryzae pv. oryzae (Xoo), the causal agent of bacterial blight, was cultured at 28°C in the dark. E. coli strains were cultured at 37°C in the dark, while the yeast strain AH109 was cultured at 30°C in the dark.

The generation of OsML1 knockout and overexpression lines was technically supported by Wuhan Boyuan Biotechnology Co., Ltd. The knockout lines, Osml1-1 and Osml1-2, were generated using the CRISPR/Cas9 genome-editing system. Specific single-guide RNAs targeting the OsML1 gene were designed and introduced into rice calli via Agrobacterium-mediated transformation. Homozygous mutants were obtained through selection and sequence verification. Osml1-1 carried a single adenine insertion, whereas Osml1-2 contained an 8-bp deletion (ΔACCAAGAA). For the overexpression lines, OsML1-OE1 and OsML1-OE2, the coding sequence of OsML1 was cloned into an overexpression vector driven by the CaMV 35S promoter, and transgenic plants were similarly obtained by Agrobacterium-mediated transformation. All T1 and subsequent generations were genotyped by PCR, and transcript levels were verified by RT-qPCR.

### Bioinformatics and structural prediction analysis

The amino acid sequences of rice OsML1, mouse MD-2, and human MD-2 were retrieved from the NCBI database and the Rice Genome Annotation Project database. Protein domain annotation was performed using SMART and the NCBI Conserved Domain Database. Three-dimensional protein structures were predicted using the AlphaFold3 online server, and the resulting models were visualized and illustrated using PyMOL. To compare the structural similarity of MD-2 proteins from different species, structural superposition was performed using TM-align, with TM-score and root-mean-square deviation (RMSD) values used as indicators of structural similarity. Multiple sequence alignment was conducted using ClustalW with default parameters, and sequence homology searches were performed using BLAST.

### RNA extraction and RT-qPCR analysis

Rice leaf samples were ground into fine powder in liquid nitrogen, and total RNA was extracted using MagZol Reagent (Magigen). Proteins and DNA were removed by treatment with BCP solution, and RNA was further purified using B10 purification columns before elution with nuclease-free water. The extracted RNA was reverse-transcribed into cDNA using the HiScript III RT SuperMix for qPCR kit (Vazyme).

RT-qPCR reactions were performed using 2× Taq Pro Universal SYBR qPCR Master Mix, gene-specific forward and reverse primers, and cDNA templates on a real-time quantitative PCR system. The amplification program was as follows: initial denaturation at 95°C for 30 s; 40 cycles of denaturation at 95°C for 5 s, annealing at 60°C for 30 s, and extension at 72°C for 30 s; followed by melting-curve analysis. OsACTIN1 (LOC_Os03g50885) was used as the internal reference gene. Relative expression levels of target genes were calculated using the 2^−ΔΔCt method. Three technical replicates were included for each sample.

### Protein expression and purification

Target gene fragments were cloned into prokaryotic expression vectors carrying either GST or His tags and transformed into competent E. coli BL21(DE3) cells. Positive clones were cultured in LB liquid medium at 37°C until the optical density at 600 nm reached 0.8. Protein expression was then induced with 0.1 mM IPTG at 16°C overnight.

Bacterial cells were harvested by centrifugation and resuspended in the corresponding binding buffer. After sonication and centrifugation, the supernatant was collected for affinity purification using magnetic beads. GST-fusion proteins were purified using GST magnetic beads and eluted with buffer containing 20 mM reduced glutathione, whereas His-fusion proteins were purified using His magnetic beads and eluted with buffer containing 250 mM imidazole. Purified proteins were stored at −80°C. Protein concentrations were determined using the Bradford assay, with bovine serum albumin (BSA) as the standard. Absorbance values at 595 nm and 470 nm were measured using a microplate reader, and protein concentrations were calculated based on the standard curve.

### LPS-binding activity assay

Recombinant expression vectors encoding full-length OsML1 without the signal peptide, OsML1^24–89, and OsML1^90–152 fragments were constructed and fused with GST and GFP tags. The constructs were transformed into E. coli for prokaryotic expression and affinity purification. Purified proteins were adjusted to a final concentration of 1 mg/mL and used freshly; they were not stored at −20°C or −80°C.

Each recombinant protein was mixed with an equal volume of LPS solution prepared in a 16-point, twofold serial dilution series. After incubation at room temperature, the binding affinity between the proteins and LPS was measured using microscale thermophoresis (MST). Binding curves were analyzed using the corresponding software, and equilibrium dissociation constants (Kd) were calculated to compare the LPS-binding capacities of different OsML1 protein fragments.

### *In vitro* bacterial binding assay

To determine the in vitro binding capacity of OsML1 to Gram-negative bacteria, particularly Xoo, the purified GST-OsML1 fusion protein was adjusted to 1 mg/mL and used freshly without freeze storage. The protein was mixed with an equal volume of Xoo suspension at an OD600 of approximately 0.8 and incubated with rotation at room temperature for 1 h.

After incubation, the mixture was centrifuged at 12,000 rpm for 10 min at 4°C to separate the supernatant and bacterial pellet. The supernatant was transferred to a new tube, while the pellet was washed three times with PBS buffer and then resuspended. Both the supernatant and resuspended pellet samples were subjected to SDS-PAGE. Empty GST protein was used as a negative control. Protein distribution was detected by Coomassie Brilliant Blue staining or Western blotting. The binding ability of OsML1 to Xoo cells was evaluated by comparing the amount of protein remaining in the supernatant before and after incubation and by determining whether the target protein was present in the bacterial pellet.

### Subcellular localization analysis

Fusion expression vectors containing full-length OsML1 or OsML1 lacking the signal peptide (OsML1-ΔSP) fused to GFP were constructed. Rice etiolated seedling protoplasts were prepared as follows. Tender stems from 7-day-old etiolated seedlings were cut into 1–2 mm segments and treated with mannitol solution for 10 min. The tissues were then incubated in cell wall digestion solution containing 1.5% Cellulase R10 and 0.75% Macerozyme R10 at 26°C and 60 rpm for 4 h.

The digestion mixture was filtered through a nylon mesh, and the filtrate was washed twice with W5 solution. Protoplasts were finally resuspended in MMg solution. For transformation, 100 μL of protoplast suspension was mixed with 10 μL of plasmid DNA at a concentration of 1,500 ng/μL. Subsequently, 110 μL of 40% PEG4000 solution was added, and the mixture was incubated at room temperature for 10 min to facilitate transformation. The protoplasts were washed twice with W5 solution and then incubated in W5 solution containing antibiotics at 26°C in the dark for 16–24 h.

Transformed protoplasts were observed using a confocal microscope. GFP fluorescence was detected under 488 nm excitation, and RFP fluorescence was detected under 561 nm excitation. The subcellular localization patterns of OsML1-GFP and OsML1-ΔSP-GFP were then analyzed.

### Measurement of reactive oxygen species burst

Rice seedlings at 7–15 days after germination, including wild-type ZH11, Osml1 knockout lines, and OsML1 overexpression lines, were used for reactive oxygen species (ROS) burst assays. Stem segments were cut into approximately 2 mm pieces and placed in 96-well plates containing sterile distilled water. The samples were kept in the dark for 10 h.

After the removal of the distilled water, 90 μL of reaction solution was added to each well. The reaction solution was prepared by adding 10 μL of luminol stock solution, giving a final concentration of 100 μM, and 20 μL of horseradish peroxidase stock solution, giving a final concentration of 20 μg/mL, to 10 mL of sterile distilled water. After addition of the reaction solution, the 96-well plates were kept in the dark for another 30–60 min to allow sufficient infiltration and stabilization of the background signal.

For the treatment group, 10 μL of LPS solution was added to each well to reach a final concentration of 100 μg/mL. For the control group, an equal volume of sterile distilled water was added. The 96-well plates were immediately placed in a microplate reader, and chemiluminescence intensity was continuously measured at room temperature. The excitation and emission wavelengths were set at 360 nm and 465 nm, respectively. Signals were recorded every 2 min for 60 min. The regulatory role of OsML1 in LPS-induced ROS burst was evaluated by comparing the peak luminescence intensity and the area under the curve among different rice lines.

### Pathogen inoculation and disease resistance phenotyping

The bacterial blight pathogen Xoo was activated by streaking on PDA solid medium and cultured at 28°C in the dark for 2 days. The activation process was repeated twice to restore bacterial vitality. Bacterial cells were washed off and resuspended in 10 mM MgCl₂ solution, and the bacterial suspension was adjusted to an OD600 of 0.6, corresponding to approximately 1 × 10⁸ CFU/mL.

Rice plants at the heading stage, including wild-type ZH11, Osml1 knockout lines, and OsML1 overexpression lines, were inoculated using the leaf-clipping method. Sterile scissors were dipped into the bacterial suspension, and rice leaves were obliquely clipped 3–5 cm from the leaf tip. Three to five leaves were inoculated per plant, with one wound generated per leaf. After inoculation, plants were maintained under high-temperature and high-humidity conditions, specifically 28°C and >80% relative humidity, with normal light exposure.

Lesion lengths were measured at 15–30 days post-inoculation and used as an indicator of rice resistance or susceptibility to bacterial blight. At least 10 leaves were inoculated for each line, and the experiment was independently repeated three times. Results are presented as mean ± standard deviation and subjected to statistical analysis.

### Metabolomic analysis

To investigate the potential role of OsML1 in rice lipid metabolism, untargeted metabolomic analysis was performed using wild-type ZH11 and Osml1 knockout mutants. Rice leaves were collected at the heading stage, with three biological replicates prepared for each genotype. Metabolite extraction, detection, and data analysis were commissioned to Tsingke Biotechnology Co., Ltd.

Untargeted metabolomic profiling was performed using a liquid chromatography–tandem mass spectrometry (LC-MS/MS) platform. Raw data were processed by peak detection, alignment, and normalization. Differential metabolites were identified using orthogonal partial least-squares discriminant analysis (OPLS-DA), with the following screening criteria: variable importance in projection (VIP) > 1.8, P < 0.05, and fold change (FC) > 2. Pathway enrichment analysis of differential metabolites was conducted using the KEGG database to identify metabolic pathways affected by OsML1 deficiency.

### Statistical analysis

All experiments included at least three independent biological replicates, each comprising at least three technical replicates. Data are presented as mean ± standard deviation (mean ± SD). Statistical analyses were performed using GraphPad Prism 9.0. Comparisons between two groups were conducted using a two-tailed Student’s t-test, whereas comparisons among multiple groups were performed using one-way analysis of variance (ANOVA) followed by

Tukey’s multiple-comparison test. Statistical significance was defined as P < 0.05, P < 0.01, and P < 0.001.

For metabolomic data, differential metabolites were screened using OPLS-DA. Metabolites with VIP > 1.8, P < 0.05, and FC > 2 were considered significantly differentially accumulated metabolites.

## Results

### Identification of an OsML1 protein with similarities to animal MD2

We previously identified rice lipid transfer protein (PITP), which has vague sequence similarities with mammalian MD-2, has a typical ML with an MD-2-related lipid-recognition domain that has LPS binding properties. MD-2 proteins take part in mammalian PAMPs signalling, and play an important role in the recognition of pathogen-associated lipid molecules, suggesting that the OsML1 protein may be directly involved in immune signal recognition. The mammalian MD-2 structural domain is involved in the recognition of bacterial lipopolysaccharide (LPS). In order to analyze if OsML1, containing the ML structural domain, has similarity with animal MD-2, we compared OsML1 with human MD-2 and mouse MD-2.

Domain structure prediction and comparison show that the three proteins all contain an ML domain with structural resemblance, indicating that OsML1 has a high degree of similarity to animal MD-2 in terms of domain composition **(Fig. 1)**.

**Figure 1.**
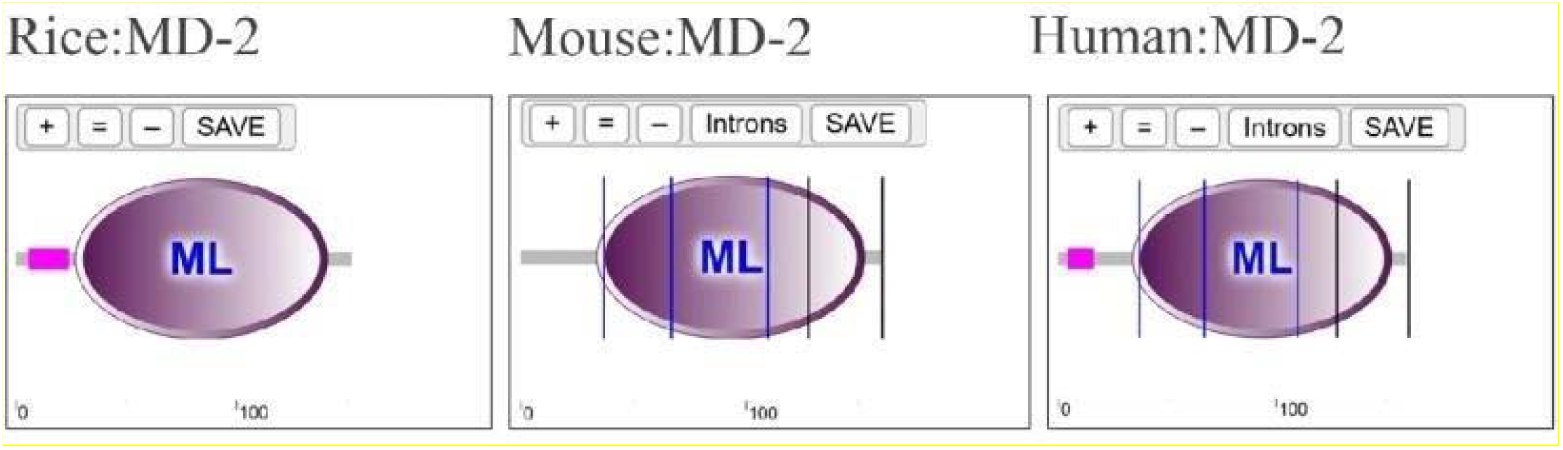
Functional domains of rice OsML1/MD-2, mouse MD2, and human MD-2 were analyzed using the SMART website. All three proteins contain a highly conserved ML domain, indicating that the ML domain, with functions in lipid recognition and signal transduction, is likely to have similar functions.

The function of MD-2 proteins in mammals and insects has been systematically analyzed, especially in relation to the toll like reseptor 4 TLR4 that plays a key role in mediating LPS recognition and immune activation. However, in plants, research on the functions of MD-2 proteins is still lacking, and there are no clear reports yet. To explore whether plant MD-2 protein and animal MD-2 have potential connections as evolutionary origin and function the amino acid sequences of MD-2 proteins from rice, mouse, and human were used for multiple sequence alignment **(Fig. 2A)**. The alignments shows that the amino acid sequences of mouse and human MD-2 are similar, while the amino acid of rice OsML1/MD-2 is less similar than the two mammalian MD-2 that has an overall sequence homology of about 36%, there is a part of the sequence that has high similarity **(Fig. 2B rice above)**. After further analysis of the predicted sequence structures, we found that the MD-2 protein of rice and mouse has most similarities towards the C-terminus **(Fig. 2B human below)**.

**Figure 2.**
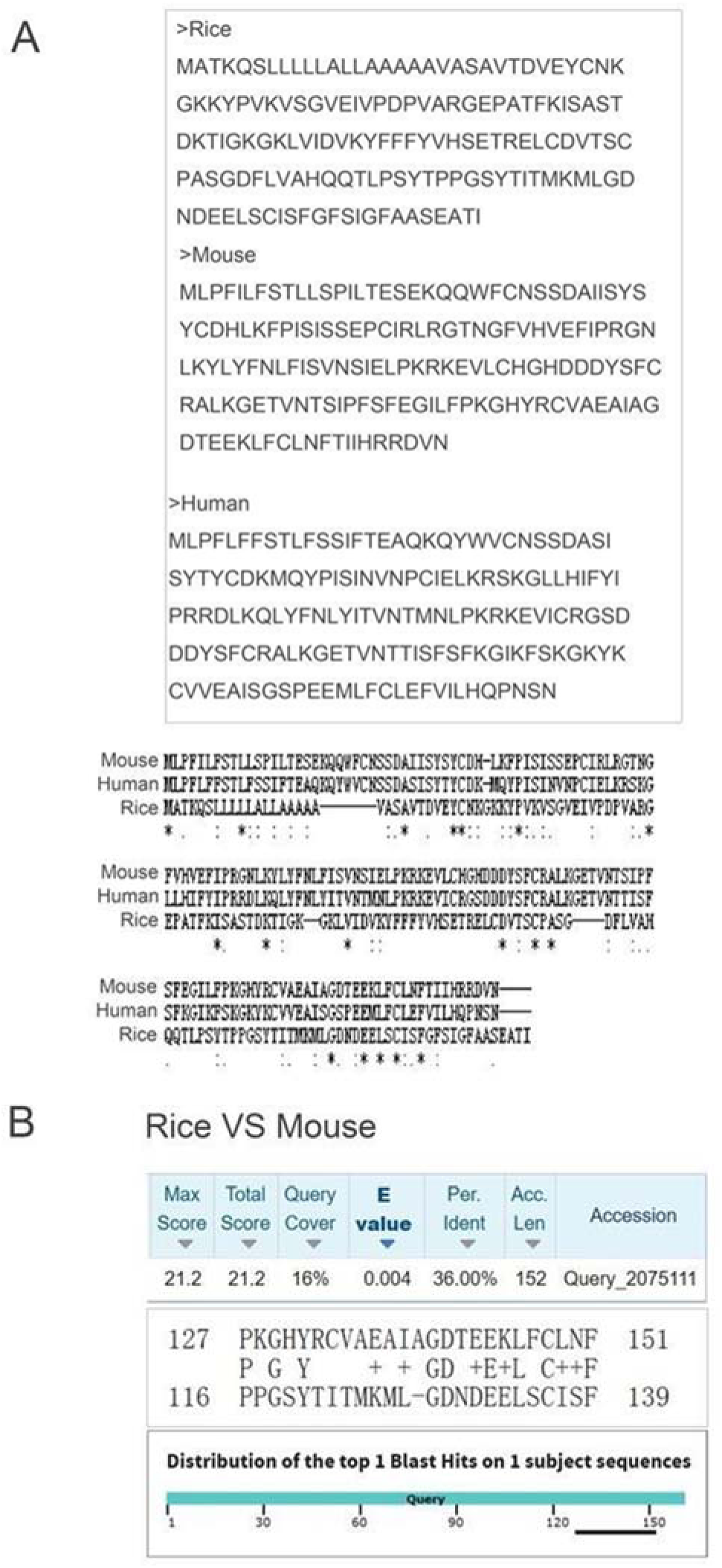
Sequence alignment and homology analysis of MD-2 across different species. (A) Full-length amino acid sequence comparison of MD-2 proteins from rice, mouse, and human shows mainly sequence similarities in the C-terminal region. (B) BLAST homology comparison between rice and mouse MD-2 proteins in the C-terminal key structural region of mouse MD-2 that is known to bind to LPS. The alignment result reveals a 36% sequence identity with 16% query coverage and an E-value of 0.004, indicating a low degree of overall homology. Despite the low conservation at the AA level, the alignment suggests that the key structural regions can be similar in structure, supporting the hypothesis that rice OsML1/MD-2 and mouse MD-2 may share equivalent LPS binding and recognition functions.

The results show that although the similarity between rice OsML1/MD-2 protein and animal MD-2 at the overall sequence level is low, there is a conservation in structural regions that recognize LPS. This may imply that OsML1/MD-2 has similar functions as mammalian MD-2 proteins or even similar evolutionary origins.

To further test OsML1/MD-2 has predicted similarities with mammalian MD-2 at the structural level with the mouse and human MD-2 LPS binding regions we used AlphaFold3 (AF3) and performed a high-precision three-dimensional folding prediction of the amino acid sequences of the three proteins and used PyMOL to visualize their predicted structures (**Fig. 3**). The modeling results show that the overall folding morphology of OsML1/MD-2 protein is similar to that of mouse and human MD-2, and all three show a typical β-barrel tructure where the interior shows hydrophobic cavities. This β-barrel structure is considered characteristic of mammalian MD-2 protein and the structural basis for lipopolysaccharide binding and activation of immune signaling pathways. It is worth noting that although OsML1/MD-2 homology with mammalian MD-2 is low in the primary sequence, its core spatial configuration shows a high structural similarity. This structural feature further suggests that OsML1/MD-2 may have similar molecular properties as animal MD-2 and thus a LPS recognition function involved in the perception and immune activation towards plant-invading Gram-negative LPS displaying bacteria.

**Figure 3.**
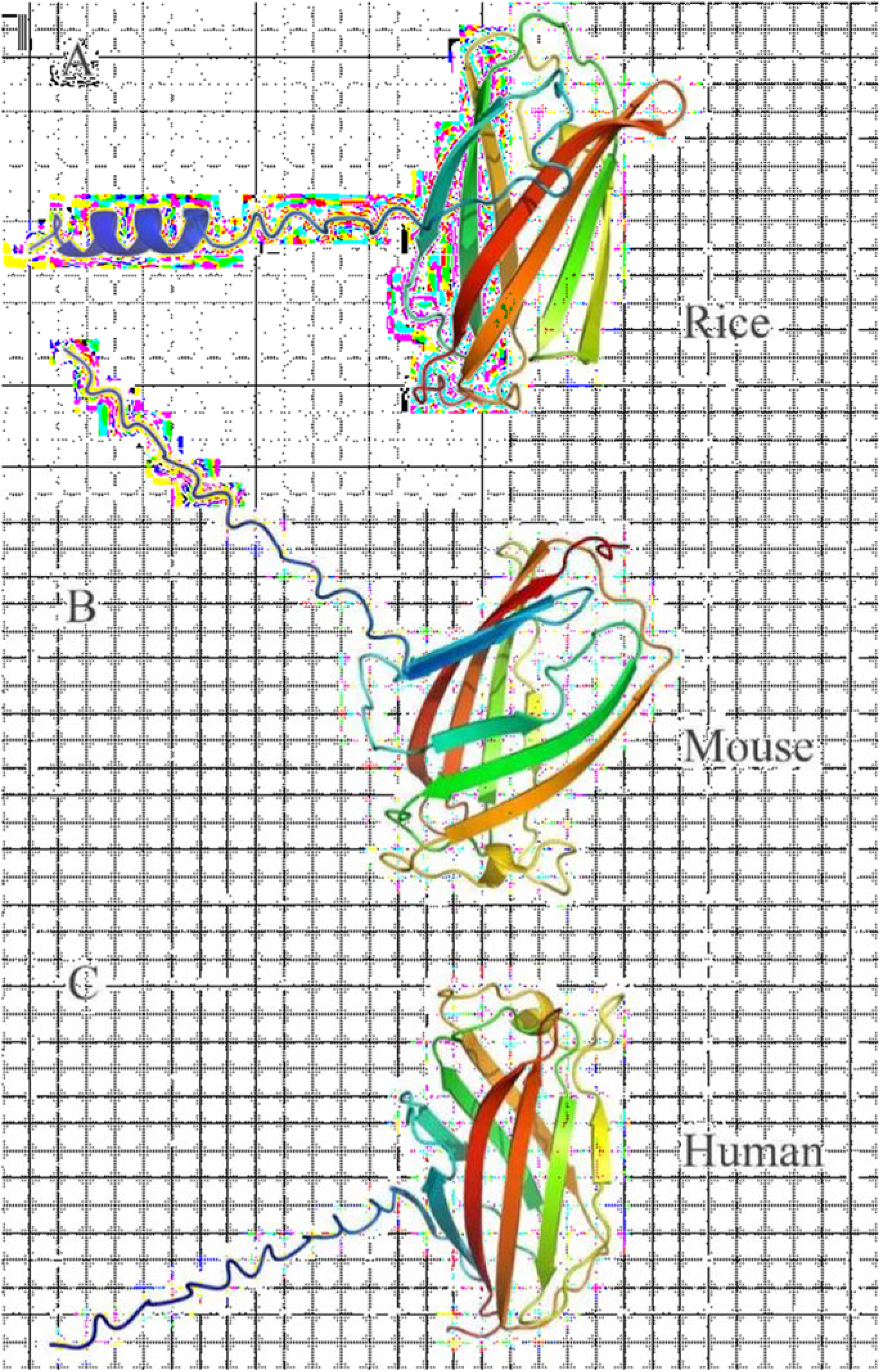
Predicted 3D structure models of MD-2 proteins of different species. (A) Predicted 3D structure of rice OsML1/MD-2 protein. (B) Predicted 3D structure of mouse MD-2 protein. (C) Predicted 3D structure of mouse human MD-2 protein.

Note that the AlphaFold3 (AF3) model still belongs to the realm of predictive analysis, and protein function conservation does not depend only on the overall structural folding. It is also affected by the retention of key functional residues and their spatial distribution. Thus, it is necessary to conduct further strict comparisons and analysis of the three-dimensional structure at the structural level. To this end, we use the structural alignment tool TM-align https://zhanggroup.org/TM-align.html) to align the three-dimensional structures of rice OsML1/MD-2, mouse MD-2, and human MD-2. Taking the OsML1/MD-2 protein as a reference mouse MD-2 protein shows similarity for a length of 121 amino acids with a TM-score of 0.57666, RMSD is 3.36 Å (**Fig. 4 A-C)**, indicating that the two proteins have similar three-dimensional conformation. According to the general standard of TM-align, a TM-score higher than 0.5 is usually judged as belonging to the same structural topological folding, suggesting that rice OsML1/MD-2 and mouse MD-2 may be structurally conserved at the functional core. Similarly, the comparison between OsML1/MD-2and human MD-2 proteins also yields similar results: the comparison length is also 121 amino acids, TM-score is 0.57440 (**Fig. 4 D-F**), with an RMSD of 3.36 Å, further supporting the conclusion.

**Figure 4.**
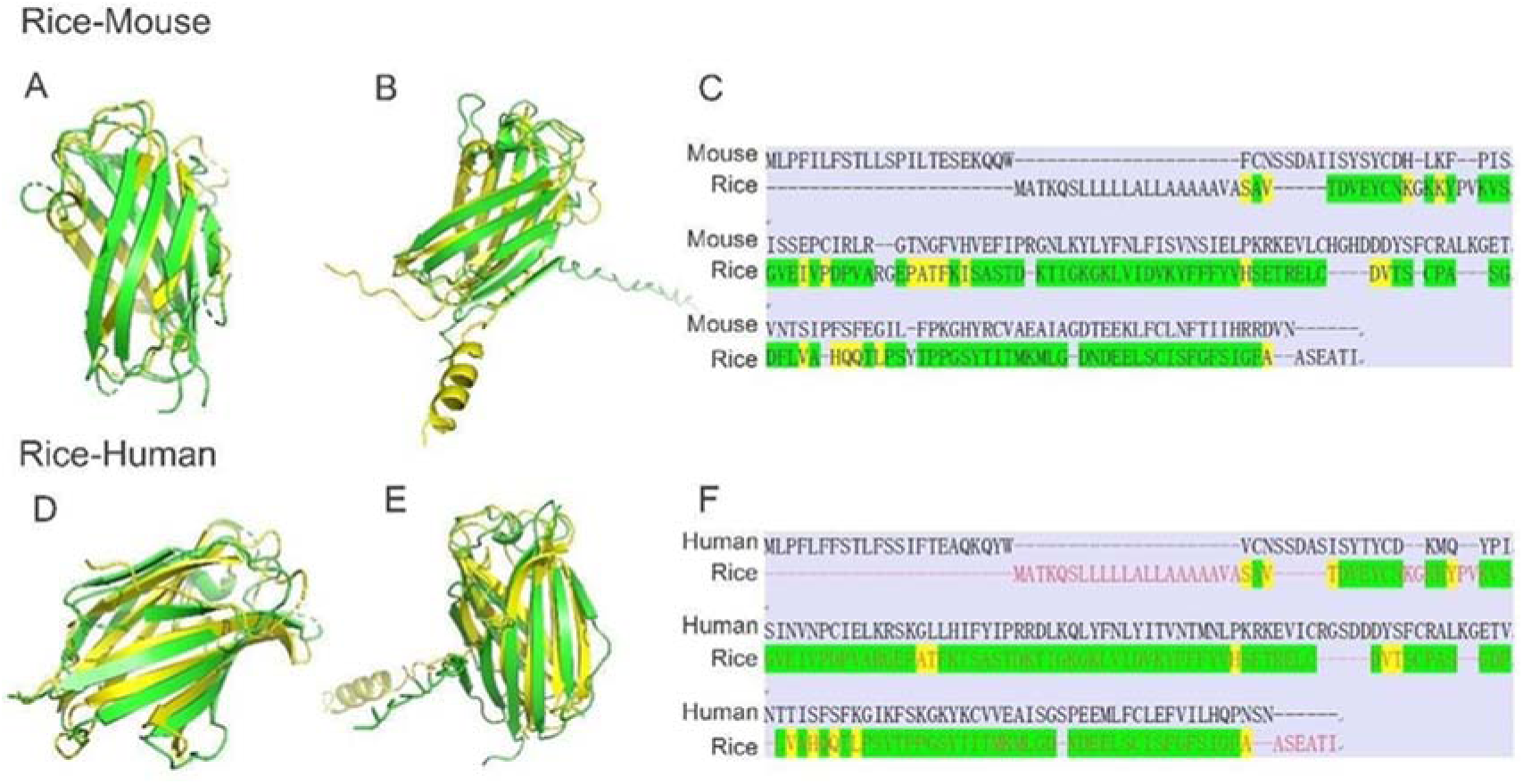
Structural alignments and sequence comparison of mammalian MD-2 and rice OsML1/MD-2. (A) Structural superimposition of the MD-2 proteins from mouse and rice in the aligned structural region. Yellow represents rice OsML1/MD-2 and green represents mouse MD-2 (B) Full-length superimposition of the rice and mouse MD-2 proteins. Yellow represents rice OsML1/MD-2, and green represents mouse MD-2. Differences are observed primarily in the N-terminal and peripheral regions. (C) Amino acid sequence alignment of rice and mouse MD-2 proteins in the aligned structural region. The partial conservation of residues within the ML domain supports potential functional convergence despite overall sequence divergence. (D) Structural superimposition of MD-2 proteins from human and rice in the aligned structural region. Yellow represents rice OsML1/MD-2 and green represents human MD-2 (E) Full-length superimposition of MD-2 proteins from rice and human. Yellow represents rice OsML1/MD-2 and green represents human MD-2 (F) Amino acid sequence alignment of rice and human MD-2 proteins. The partial conservation of residues within the ML domain supports potential functional convergence despite overall sequence divergence.

OsML2 is very similar in sequence to OsML1, indicating that the latter can have evolved from a gene multiplication of OsML1 (**Fig. 5A**), but it is OsML1 that has a typical β-folded core structure, as MD-2 proteins have (**Fig. 5B**), which has a main role and is upregulated in response both to abiotic and biotic stresses linked to plant infection and particularly to bacterial Xoo infection (**Fig. 5C-H**)

**Figure 5.**
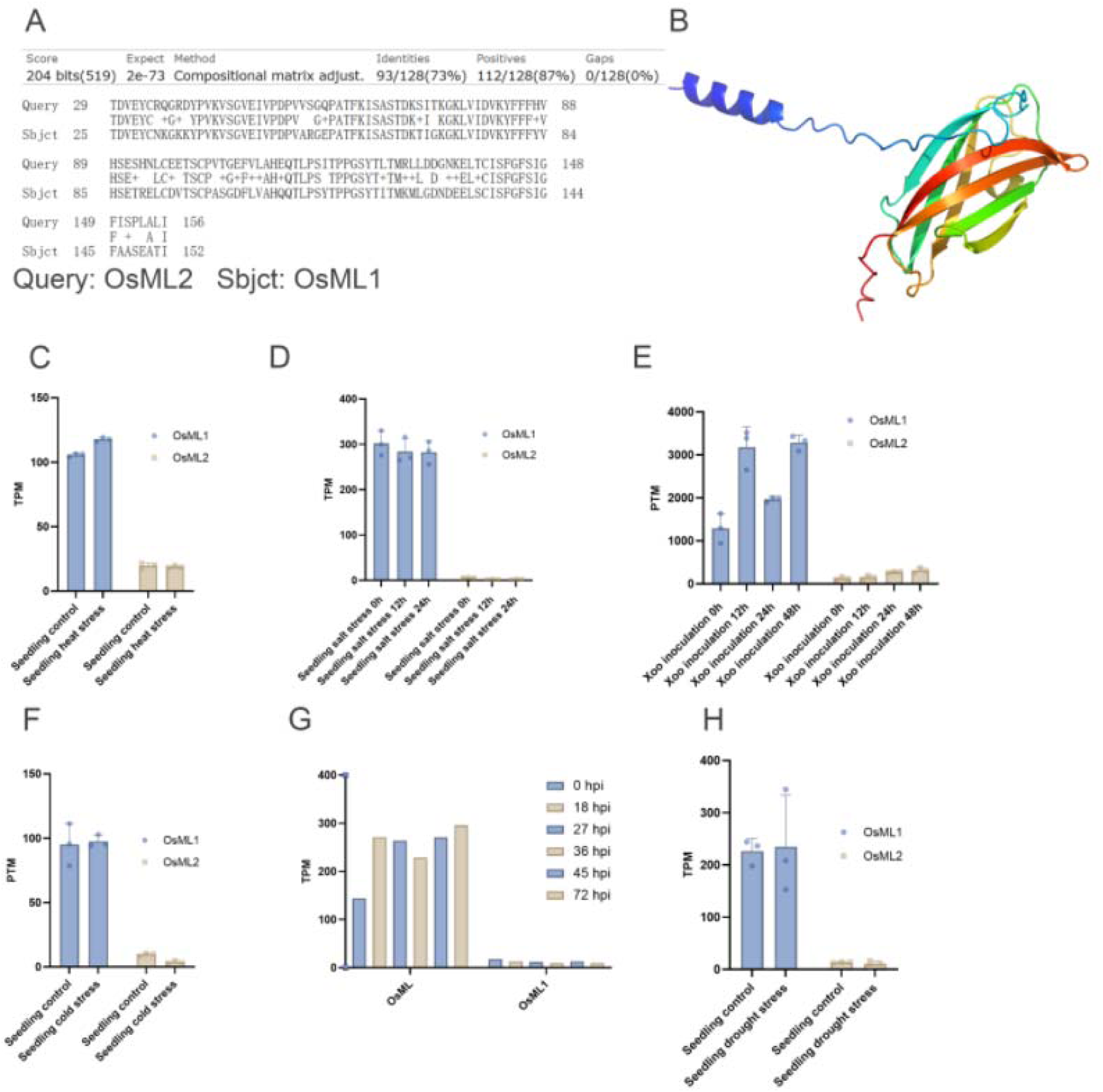
Structural and expression analysis of rice OsML1 and OsML2. (A) OsML1 and OsML2 amino acid sequence alignment. The alignment results show that the proteins have high similarity at the core ML domain similarity, suggesting conserved functional structures, but there are some larger sequence differences in the N-terminus. (B) Three-dimensional OsML1 structural prediction diagram. It shows a typical β-folded core structure, which conforms to the structural characteristics of the ML family proteins. (C-F) Comparison of TPM expression levels of OsML1 and OsML2 under different stress treatments: (C) cold stress; (D) heat stress; (E) dry drought stress; (F) salt stress. The results show that the expression of OsML1 under various abiotic stress conditions is higher than that of OsML2, suggesting that OsML1 may play a more active role in environmental stress response. (G) OsML1 and OsML2 in rice blast fungus show different FPKM expression changes over time. OsML1 is upregulated in the early stage of infection and may participate in the initial stages of pathogen recognition. (H) FPKM expression changes of OsML1 and OsML2 at different time points after infection with paddy rice leaf blight bacteria. OsML1 is significantly highly expressed at multiple time points, suggesting that it plays a more important role in the immune response under bacterial stress.

The structural domain comparison results were visually compared using PyMOL visualization analysis. This analysis shows that OsML1/MD-2 and mammalian MD-2 not only present a high degree of consistency in the overall folding morphology but also show a strong spatial overlap. This is an example of proteins with high three-dimensional structural conservation, even with low sequence homology, which is an evolutionary characteristic of distant homologs or convergent evolution at the sequence level differentiation, but the key functional domains of the structure are retained or formed due to selective constraints.

In summary, OsML1/MD-2 protein is highly similar to the mammalian MD-2 when it comes to the essential spatial folding structure, suggesting that it may have a similar function in lipid recognition and innate immunity activation that has not been previously demonstrated in plants.

### OsML1 is secreted and does not have an antibacterial function in vitro

In mammals, MD-2 proteins are mainly located in the cytoplasm and are also secreted extracellularly. They co-activate the immune signal mediated by the cell surface toll-like receptor 4 (TLR4) by specifically binding to lipopolysaccharide, thereby initiating a typical first-day innate immune response.

In order test a possible secretion of OsML1, we constructed an expression vector of OsML1 and transformed it into a yeast system for expression. The yeast secretion experiment results show that the complete OsML1 protein can be effectively secreted from yeast cells, its secretion is similar to the positive secretion control (**Fig. 6A**), indicating that OsML1 is cotranslationally secreted. It is worth noting that when the DNA sequence encoding the predicted OsML1’s N-terminal signal peptide sequence was removed, the secretion of the protein almost completely stopped, indicating that the signal peptide is necessary for the OsML1 protein to enter the normal secretory pathway.

**Figure 6.**
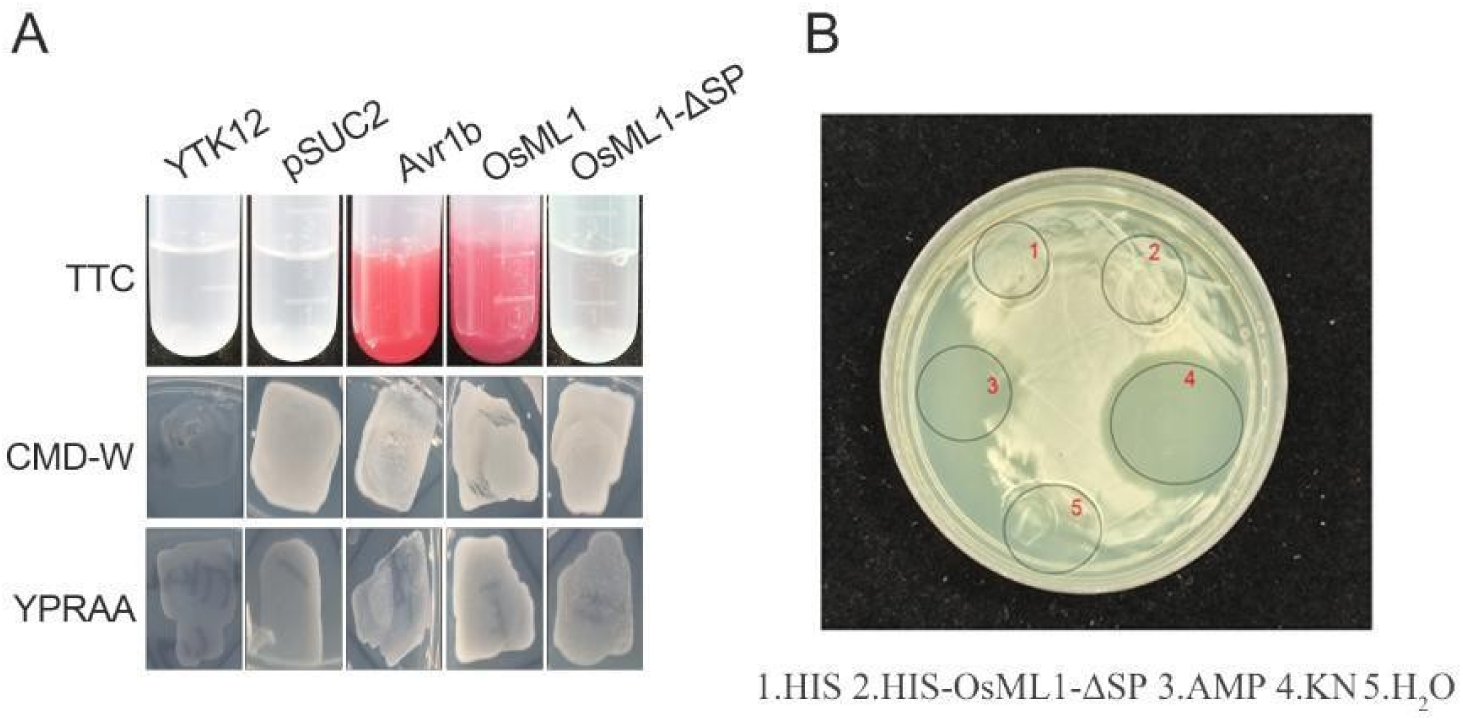
Signal peptide activity and in vitro antimicrobial analysis of rice OsML1. (A) Signal peptide secretion activity detection of OsML1. YTK12 and pSUC2 are negative controls, and Avr1b is a positive control. (B) In vitro antibacterial activity detection of OsML1 protein. The antibacterial ability of different protein solutions against target pathogens was analyzed by the colony diffusion method. (1) HIS empty carrier protein (negative control), (2) HIS-OsML1-ΔSP protein (HIS-OsML1 with signal peptide removed protein), (3) 100 ng/mL AMP (antibiotic control), (4) for 50 ng/mL KN (antibiotic control), (5) H₂O (negative control). CMD-W and YPRAA show yeast cell growth on the respective medium

Furthermore, in order to evaluate whether OsML1 has a direct antibacterial activity, since it could have that due to its resemblance with MD-2 and bind to Gram-negative bacterial LPSs in an antibody-like fashion, we constructed a GST-tagged fusion expression in the KN vector and successfully purified OsML1 proteins *in vitro*. We applied the purified protein to the surface of *E. coli* grown in LB medium without antibiotics, with an antimicrobial peptide (AMP) tagged fusion expression vector, and successfully purified OsML1 proteins in vitro. As controls, we used the purified and empty KN vector and H_2_O. No obvious zone of inhibition was observed (**Fig. 6B**), suggesting that OsML1 has no direct antibacterial activity.

OsML1 has lipopolysaccharide binding activity and can bind to *Xanthomonas oryzae* pv. oryzae in vitro

To investigate the LPS binding ability of the OsML1 MD2 domain in rice we constructed vectors encoding several different OsML1 protein fragments including, the full-length OsML1 protein without signal peptide (OsML1-ΔSP), OsML1 24–89 containing amino acid segment (OsML1 24–89), and segment OsML90–152 containing amino acid segment (OsML1 90–152), and fused them to the GST-GFP vector and expressed them in *E. coli*. After purification, the expressed proteins were used for in vitro binding experiments. The results showed that the full-length OsML1 protein has LPS binding ability. It shows a strong affinity, and its equilibrium dissociation constant (Kd) that significantly reduced with increasing concentration, indicating that OsML1 and LPS is efficiently and stable binding (**Fig. 7A**). Further comparison of LPS binding of different regional fragments showed that OsML1 24–89 has a similar binding ability as the full-length protein although its affinity is slightly higher than full length OsML1 while the OsML1 90–152 segment shows hardly any binding activity to LPS (**Fig. 7A**). These results clearly points out that the binding of OsML1 to LPS is mainly N-terminal region mediated.

**Figure 7.**
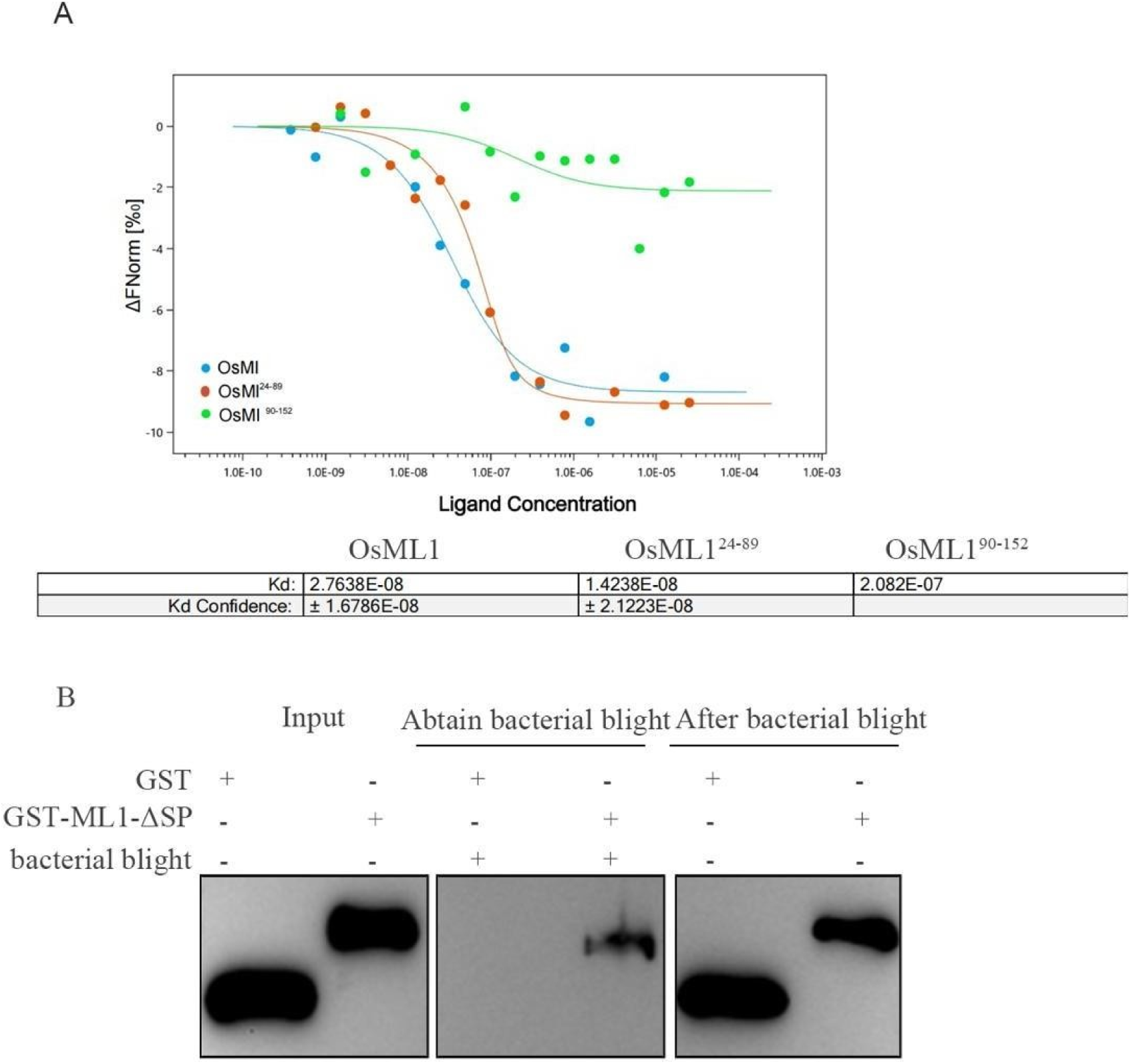
Affinity analysis of OsML1 for binding to LPS and an *in vitro* binding analysis to Xoo. (A) Fluorescence quenching analysis of the binding ability of OsML1 to LPS. OsML1 protein fragments of different lengths (full-length OsML1, OsML1 24-89, and OsML1 90-152) were used for binding curve determination. The horizontal coordinate shows ligand (LPS) concentration, and the vertical coordinate shows the change in fluorescence intensity. The results showed that OsML1 24-89 has the strongest binding ability (Kd 1.42 E-08 M), slightly stronger than full-length OsML1 (Kd = 2.76E-08M), while the C-terminal fragment OsML1 90-152 binding ability almost disappeared. (B) Protein binding experiment to detect the change in the binding ability of OsML1 to samples before and after Xoo treatment.

But can OsML1 bind not just LPSs but also intact Gram-negative bacteria *in vitro*, and then especially the rice bacterial blight pathogen *Xanthomonas oryzae pv. oryzae* (Xoo)? To test this OsML1 protein solution was co-incubated with Xoo bacterial cells, and the protein binding to the cells was analyzed by centrifugation to separate the non-bacteria-binding OsML proteins left in the supernatant from those that bind to the bacterial cells and co-precipitate with the cells in the pellet.

The results showed that the amount of OsML1 protein in the supernatant did not change significantly before and after treatment with only the empty GST protein vector used as a control or when OsML1 lacked the signal peptide (OsML1-ΔSP). However, when incubated with Xoo, the OsML1 proteins left in the supernatant were markedly reduced, indicating that OsML1 can interact with and bind to intact Xoo cell surface structures (**Fig. 7B**).

In summary, our experimental results show that the OsML1 protein not only can bind LPS in vitro, but its main LPS-binding region is in the N-terminal part of the protein, and it can bind directly to Xoo cells.

### OsML1 subcellular localization in rice cells

To verify that the OsML1 signal peptide determines its predicted localization in rice cells, we constructed a full-length OsML1-GFP fusion expression vector with a green fluorescent protein marker as well as an OsML1-ΔSP-GFP fluorescent fusion expression vector with the signal peptide removed. Subsequently, the two vectors were introduced into rice protoplasts by PEG-mediated transient transformation and incubated on a horizontal shaker under light for 16 hours. Confocal fluorescence microscopy was used to observe the GFP-labelled proteins and their subcellular localization.

The experimental results showed that the full-length OsML1-GFP protein was mainly located in the cytoplasm but also showed typical membrane association characteristics, while the OsML1-ΔSP-GFP protein without the signal peptide showed a uniform fluorescent signal distributed throughout the protoplast cytoplasm, thus lacking obvious membrane localization (**Fig. 8**)

**Figure 8.**
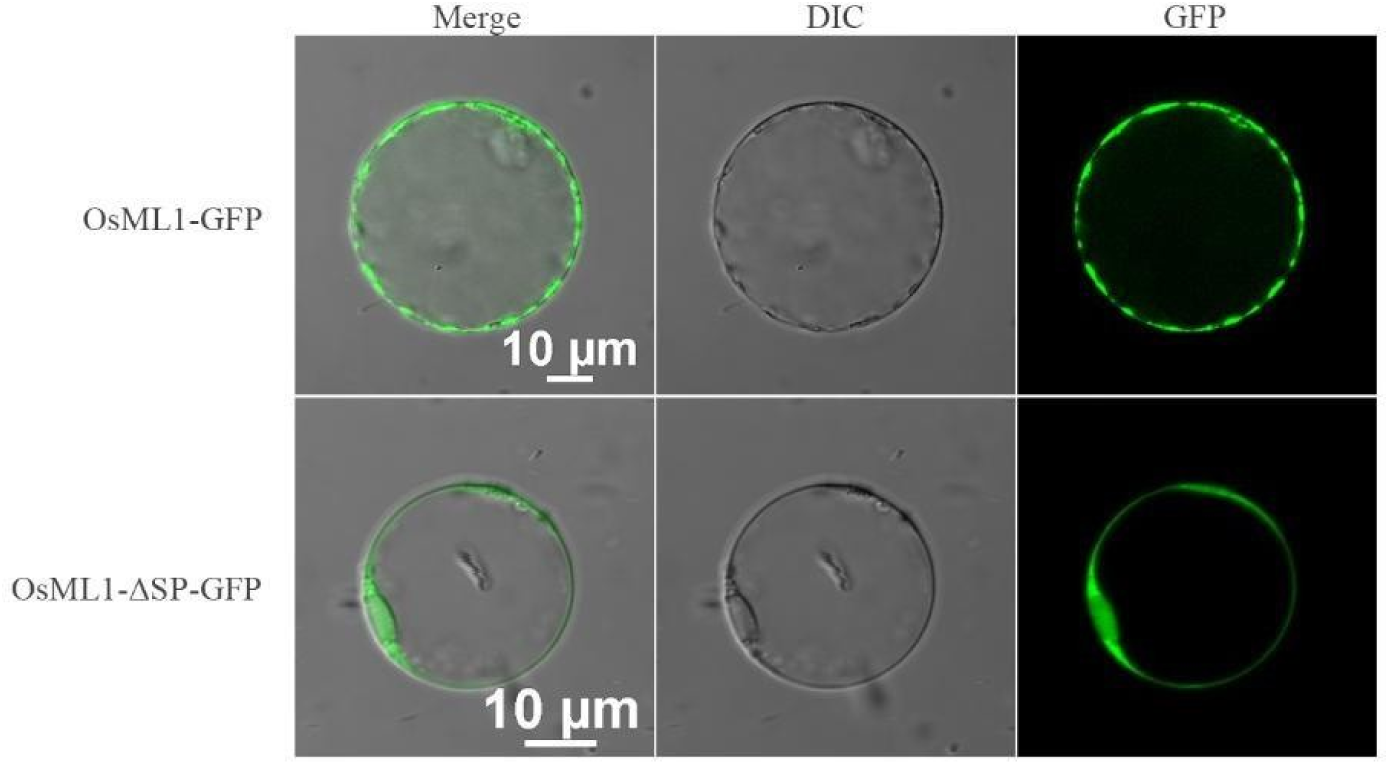
Subcellular localization of OsML1-GFP and OsML1-ΔSP-GFP in rice protoplasts, showing a distribution pattern of OsML1-GFP that is typical for a membrane-localized protein.

The above results indicate that the native signal peptide of the OsML1 protein can guide the protein to the secretory pathway, so it may appear at the plant cell membrane.

### OsML1 responds to lipopolysaccharide signals and positively regulates rice resistance towards bacterial blight

We constructed OsML1 knockout and overexpression strains, and performed disease resistance phenotype analysis using CRISPR/Cas9 gene editing technology to systematically explore the function of OsML1 in rice immune response. Two homozygous mutant lines were obtained in the T1 generation of plants. The first one is a single A base insertion mutation (named *Osml1-1*), and the other is an 8 bp deletion mutation (ΔACCAAGAA, named *Osml1-2*) (**Fig. 9A**). In addition, we constructed an overexpression line driven by the 35S promoter of OsML1 to enhance OsML1 expression. We used RT-qPCR for verification of the relative expression levels of the different constructs in plant tissues compared to *OsML1* expression in the background (ZH11). The results showed that the transcript expression of *Osml1-1* and *Osml1-2* was significantly downregulated, while *OsML1* was significantly upregulated in the overexpression lines **(Fig. 9B)**.

**Figure 9.**
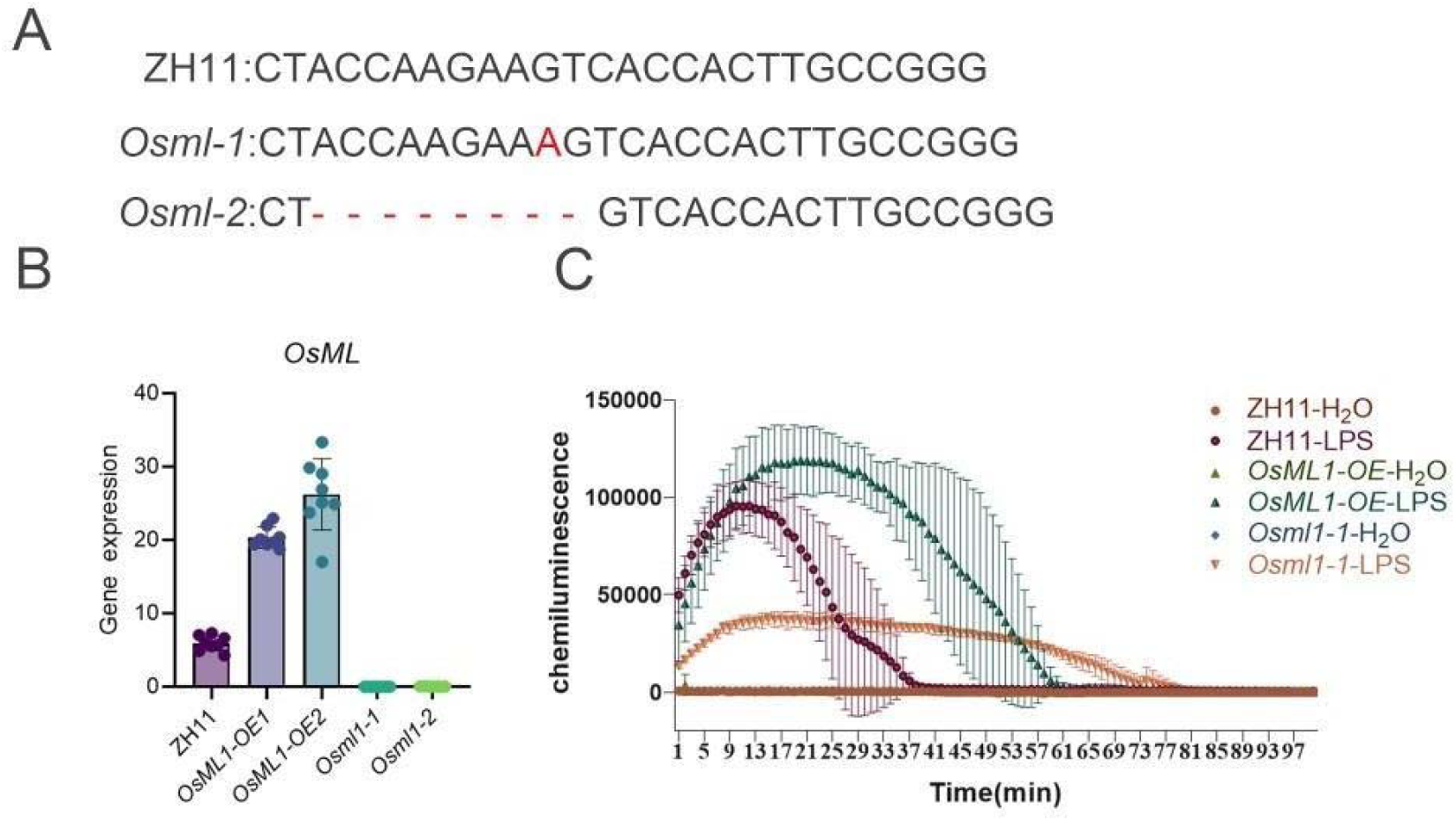
OsML1 regulates LPS-induced ROS burst levels. (A) OsML1 Gene editing sites and mutation types. (B) Analysis of transcriptional expression levels of *OsML1* genes in Different Plants. *OsML1* is significantly upregulated in overexpression lines (*OsML-OE*), while its expression signal is almost absent in mutants (*Osml1*). (C) Analysis of ROS burst levels induced by LPS in different plant lines. ROS burst in *Osml1-1* plants with the G to A mutation in the *OsML1* gene was significantly weaker compared to wild-type ZH11 plants, while ROS accumulation was notably enhanced in *OsML-OE* plants, demonstrating that OsML1 proteins are positively involved in creating LPS-induced ROS burst levels.

Given that LPS is a PAMP that can induce rapid ROS immune responses, especially in plants, we compared the ROS accumulation levels in the different plant lines after LPS treatment.The results showed that the ROS burst of *Osml1-1* plants was significantly weakened, while the ROS accumulation in OsML1 overexpressing plants was significantly increased compared to the background (ZH11) **(Fig. 9C)**. These results indicate that OsML1 is involved in the plant’s perception and response to LPS and is an important regulator of LPS-induced ROS accumulation characteristic of plant immune responses.

We designed a leaf-injury inoculation experiment on detached rice leaves to test if OsML1 affects rice plant resistance to bacterial blight (Xoo) inoculation and to evaluate the effects of the *OsML1* deletion and expression level on rice bacterial blight resistance. Rice variety ZH11 was used as the wild-type control, and two OsML1 gene knockout plant lines (Osml1-1 and Osml1-2) were used, as well as two overexpression lines (OsML1-OE1 and OsML1-OE2). On day 10 after Xoo inoculation of the leaves, lesion lengths were measured and statistically analysed to evaluate disease severity. The results showed that compared with wild-type ZH11, the lesion lengths of Osml1-1 and Osml1-2 were significantly longer, indicating that the Xoo pathogen spread inside the leaves was markedly increased (**Fig. 10**). This indicates that OsML1 deletion reduces paddy rice resistance to bacterial blight by weakening the plant’s defense responses. In line with this, OsML1 overexpressing strain OsML1-OE-1 and OsML1-OE-2 showed significantly shortened lesion lengths compared to ZH11, indicating a stronger disease resistance (**Fig. 10**), suggesting that overexpression of OsML1 can enhance rice resistance to bacterial blight.

**Figure 10.**
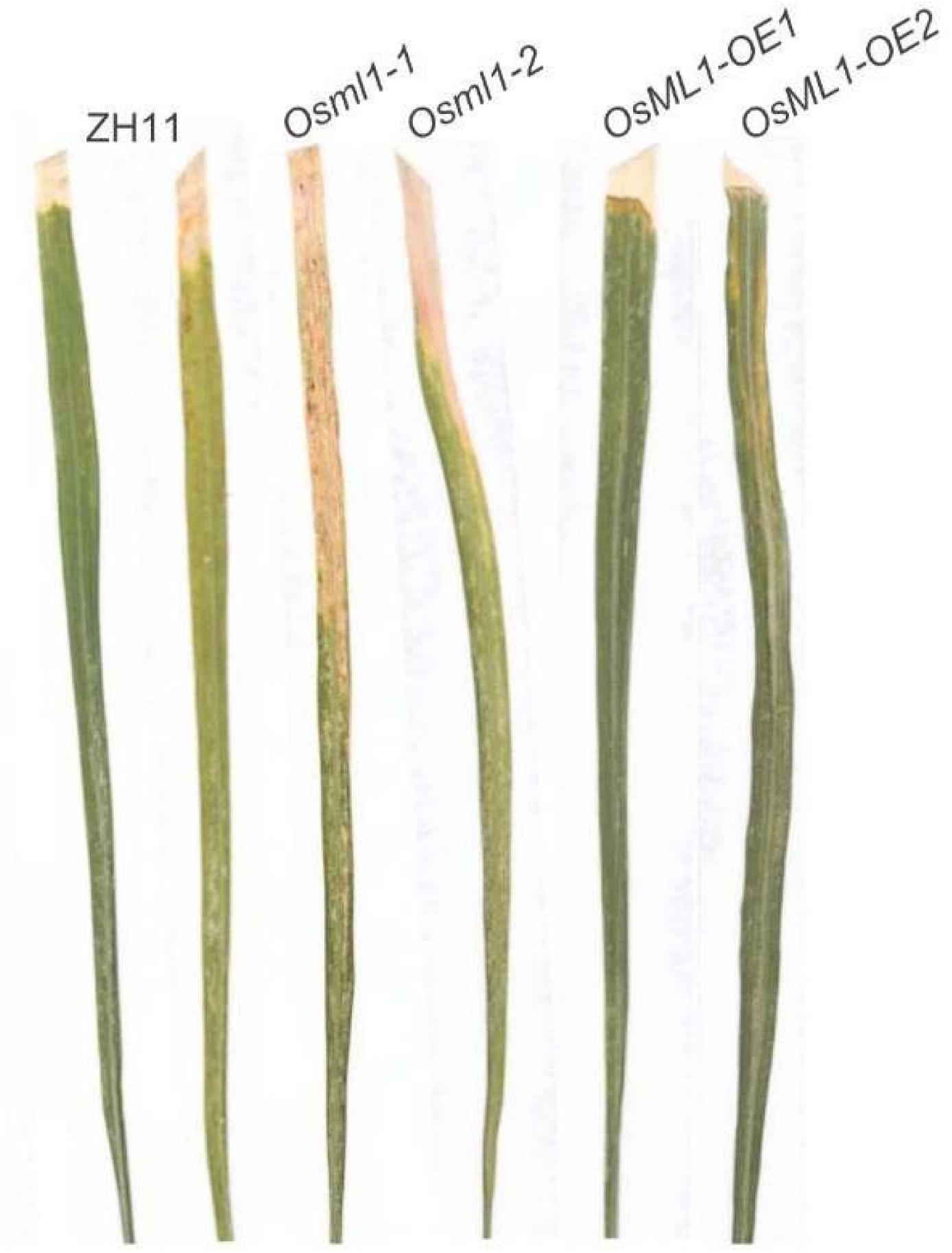
Xoo disease development on different plant lines. Rice plants with the *OsML1* gene mutated *Osml1-1* and *Osml1-2* show increased disease symptoms compared to the background ZH11 when leaves were challenged with Xoo, while rice plant lines overexpressing *OsML1* (*OsML1-OE1* and *OsML1-OE2*) show decreased disease symptoms when similarly challenged.

Taken together with the previous results on the binding ability of OsML1 to LPS, the different plant lines’ ROS responses to LPS addition provide key experimental evidence that OsML1 functions as a plant lipopolysaccharide (LPS) recognition factor, or receptor protein. OsML1 mediates the LPS-triggered immune activation and pathogen responses involved in plants (at least grasses) and is a likely candidate for a protein directly involved in LPS-binding and perception, triggering plant innate immunity towards Gram-negative pathogenic bacteria invading their tissues.

### OsML1 is widely involved in rice metabolic processes

Our previous study [2] has shown that OsML1 is a protein with lipid-binding ability, suggesting that it may be involved in plant metabolism as a lipid transporter. We carried out a non-targeted metabolomics analysis of wild-type rice and OsML1 gene deletion mutant materials to gain more insight.

Analysis of the lipid metabolomic data detected a total of 690 metabolites that were significantly differently expressed in the ZH11 background compared to in *Osml1* mutant plants. Significantly upregulated in *Osml1* mutant plants were 453, while 237 metabolites were significantly down-regulated (**Fig. 11A**), indicating that the deletion of *OsML1* significantly reshaped the lipid metabolic regulatory network in rice.

**Figure 11.**
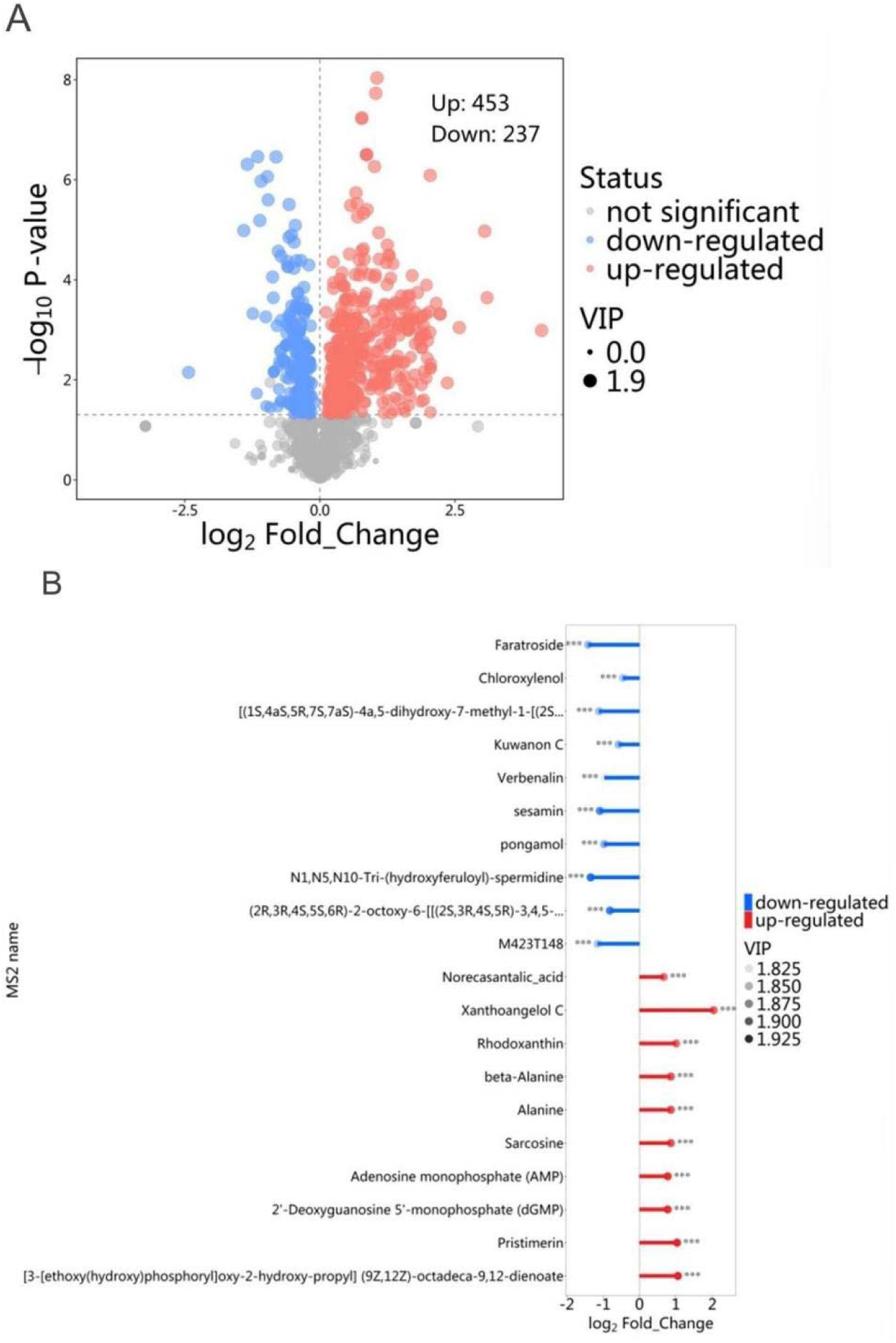
Metabolomic effects in Osml1 plants compared to background plants. (A) Volcano plot of differentially accumulated metabolites in *Osml1* plants compared to wild-type plants (ZH11) Note: This figure shows differential accumulated metabolites at the metabolomic level between the Osml1 mutant and the wild-type ZH11. X axis Represents log 2 Fold Change. The Y-axis is -log 10 (P-value), and is used to show significance and fold change of differentially accumulated metabolites. Red Colored dots represent metabolites with increased abundance (453) and upregulated synthesis, blue dots represent metabolites with decreased (237 species) downregulated synthesis, and gray dots represent metabolites with significantly different changes in synthesis. (B) Analysis of significantly different metabolites between the Osml1 mutant and wild-type ZH11. Note: This figure displays the top significantly differentially accumulated metabolites (DAMs) screened between the Osml1 mutant and wild-type ZH11 with criteria of VIP values > 1.8, P < 0.05, and a Fold Change> 2. Red bars represent up-regulated metabolites, blue bars represent down-regulated metabolites. The horizontal axis shows log2 Fold Change. The VIP value of each metabolite is represented by the dot size, and the statistical significance is indicated by asterisks where *=P<0.05, **=P<0.01,***=P<0.001.

To further analyze the specific characteristics of these metabolic differences between the *Osml1* mutant and the ZH11, significantly changed metabolites were screened, to draw a lollipop heatmap (**Fig. 11B**), with FC representing the magnitude of changes in metabolite expression, and combine that with VIP values (Variable Importance in Projection) to evaluate the metabolite importance in the found differences.

Beta-alanine, which is related to increased cellular oxidative stress were significantly upregulated in *Osml1* knockout plants. And sarcosine, which is an intermediate in glycine metabolism, is likewise upregulated. These two upregulations suggest that OsML1 deficiency may affect stress responses by influencing the synthesis of amino acids and thereby redox homeostasis.

The content of the monophosphates AMP and dGMP, related to energy metabolism and the low-energy form of both ATP and GTP, also increased significantly in the knockout plants, indicating that deleting OsML1 may disrupt the energy metabolism within the cell, lowering cell activity.

Among the down-regulated metabolites that decreased significantly in the knockout plants, we found compounds with known biological activity, lignan (such as Sesamin) and flavonoids (such as Kuwanon C). These secondary metabolites are known to play key roles in plant stress response, to adverse conditions, in antioxidant defence, and immune signal regulation. The decline in these compounds suggests that the OsML1 deletion may weaken the chemical defenses of rice.

In addition, metabolites like chloroxylenol that are known to have broad-spectrum antibacterial activity were also significantly down-regulated in the OsML1 knockout plants, further suggesting a positive effect of OsML1 in regulating both rice leaf water potential and antibacterial metabolite production as part of the immune response.

A KEGG pathway enrichment analysis was performed to find significantly differently expressed metabolites. The analysis results show that metabolites with changed expression are mainly enriched for amino acid metabolism, carbohydrate metabolism, energy metabolism, nucleotide metabolism, membrane transport, and translation-related pathways (**Fig. 12A)**.

**Figure 12.**
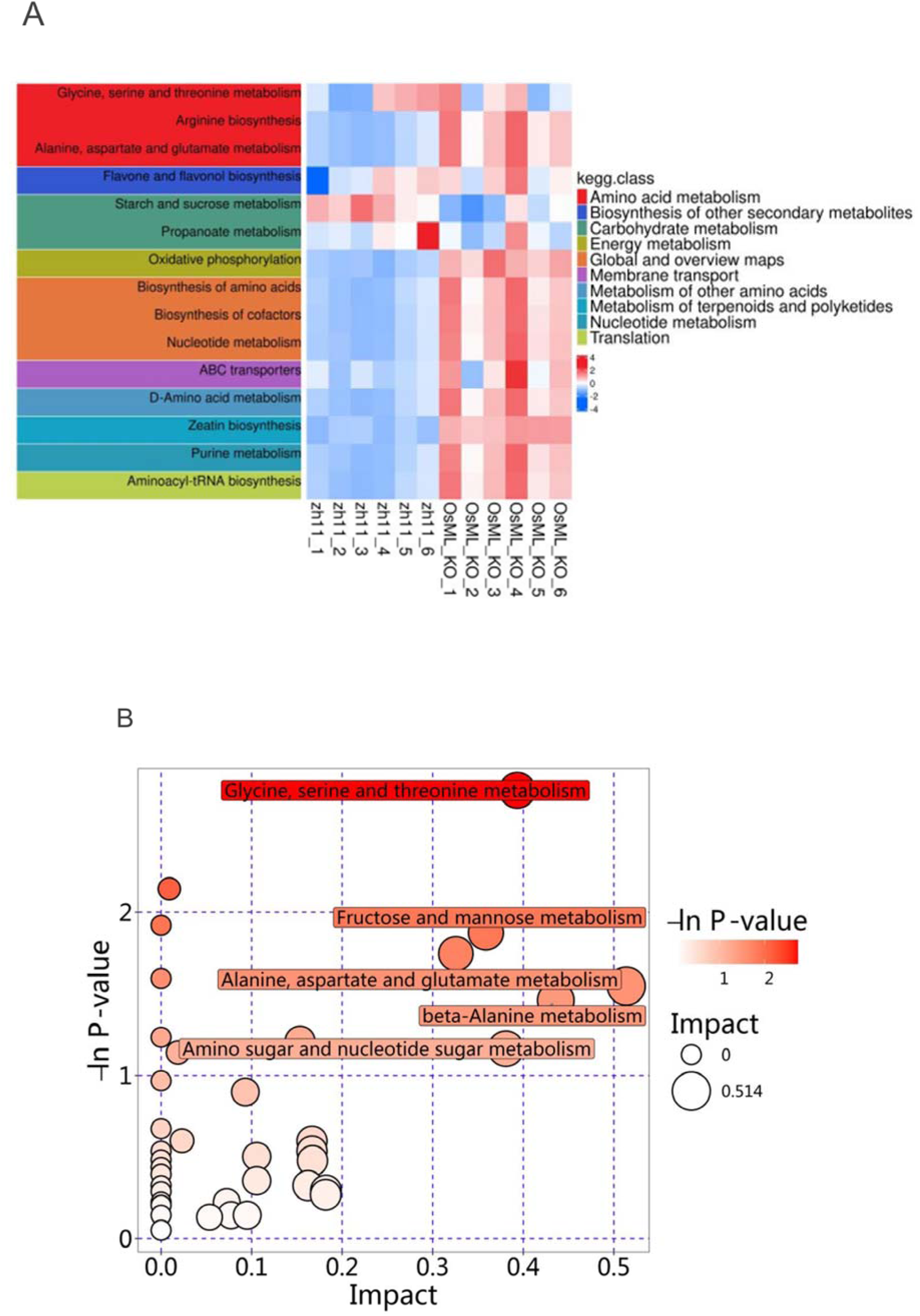
Enrichment heatmap and pathway/impact analyses of methabolites. (A) KEGG-based enrichment heatmap of differentially regulated metabolic pathways The figure shows the enrichment of metabolic pathways in different replicate *Osml1* mutant plants and wild-type plant (ZH11) samples. The X-axis shows sample numbers. The Y-axis shows the significantly enriched KEGG metabolic pathways. Blue to red indicates the degree of enrichment (z-score), blue indicates relative enrichment, and red indicates relative depletion. (B) Pathway enrichment and impact analysis of differentially accumulated metabolites in *Osml1* mutant plants compared to wild-type (ZH11) plants The figure shows the metabolic differences obtained through metabolic pathway analysis (MetaboAnalyst) of different KEGG pathways. The X axis shows pathway impact factor values (Pathway Impact), that represents the change in the metabolic pathway; the Y axis is the pathway enrichment significance (-log10 P-value). The larger the value, the more significant the pathway enrichment. The bubble color from light to dark indicates the significance of the P value, and the bubble size represents the size of the impact factor.

Thus, the OsML1 deletion has a wide range of effects and impacts rice’s primary metabolism and molecular functions.

Further, visualization enrichment analysis (**Fig. 12B)** revealed that glycine, serine, and threonine metabolism pathways showed the most significant changes in the Osml1 knockout plants with the lowest p value and the highest pathway impact value (Impact), suggesting that this metabolic pathway is the most significantly enriched among the functional pathways affected by the knockout. That pathway is widely involved in nitrogen metabolism, protein synthesis, signal transduction, antioxidant defense, and other important life activities, suggesting that this pathway, which has a profound impact on the stress response and metabolic homeostasis of rice, may be regulated by OsML1.

In addition, fructose and mannose metabolism, alanine, aspartate, and glutamate metabolism, β-alanine metabolism, as well as amino sugar and nucleotide sugar metabolism, and other pathways also showed significant differences. These pathways involve cell energy and multiple core cellular processes such as nutrient assimilation, amino acid transport, antioxidant reactions, stress responses, cell wall modifications, and sugar metabolism.

Significant upregulation of β-alanine (Beta-Alanine) and sarcosine (Sarcosine) was found in the mutants that may reflect a compensatory response of plants to oxidative stress. Similarly, upregulation of lignans (such as Sesamin) and flavonoids (such as Kuwanon C) may show a weakened plant’s stress resistance and immune defense ability, since these compounds are generally closely related to antioxidant activity, and are necessary for efficient pathogen defense.

Adding to this, the significant down-regulation of antibacterial-related metabolites (such as Chloroxylenol) further indicates that OsML1 deletion may also weaken the plant’s chemical defense barrier against pathogens, thereby affecting its overall disease resistance.

Taken together, these findings suggest that OsML1 may act as a link between primary metabolism and key nodes in plant immune responses and play a central role in rice’s response to biological stresses.

## Discussion

### OsML1 has typical MD-2 structural features and LPS binding ability

Our functional prediction of OsML1 starts from its sequence and structural analysis using SMART, NCBI, CDD, and AlphaFold3 tools to perform domain annotation and three-dimensional modeling of the OsML1 protein. We found that OsML1 contains a typical ML domain with a β-barrel folding structure similar to that of mammalian MD-2 protein [7,8]. Although the homology of the MD-2 protein between paddy rice and human or mouse at the primary sequence level is low, only about 30%. However, TM-align structural alignment shows that their three-dimensional structures are highly similar, the TM-score is 0.57, exceeding the 0.5 threshold. The topological structure similarity suggests that they may have functionally comparable spatial folding and conformation (**Fig. 4**).

Further experiments confirmed that OsML1 protein can bind lipopolysaccharide with high affinity in vitro, and its Kd value is as low as on the nM level, indicating that OsML1 may directly bind to and sense LPS molecules (Fig. ??). We further analyzed OsML1 protein by fragment cloning and tested the binding ability of different modified structural domains, and it was found that the lipopolysaccharide binding region is mainly concentrated in the amino acid region 24-89. (**Fig. 7)**This region is important for forming β-folded structures that may be the basis for the formation of a hydrophobic cavity. In contrast, the amino acid region 90-152 did not show significant binding ability, confirming that the lipopolysaccharide binding function has a region specificity (**Fig. 7**).

Lipopolysaccharide is an important component of the cell wall of Gram-negative bacteria and is a typical PAMP in plant basic immune responses [4,5]. OsML1 can bind to lipopolysaccharide in vitro and bind to the Gram-negative rice pathogen *Xanthomonas oryzae pv. oryzae*. The binding of bacteria suggests OsML1 may exist as a plant cell surface receptor for plant immune recognition. We have further shown that OsML1 can sense exogenous LPS and also positively respond to signals inducing a ROS burst, since the LPS induced ROS burst was significantly weakened in *Osml1* mutant plant, while *OsML1* overexpression significantly enhanced the ROS response, indicating that OsML1 can induce LPS triggered PTI-responses. *Xanthomonas oryzae pv. oryzae* (Xoo) leaf cutting inoculation experiments further showed that knockout strains (Osml1-1 and Osml1-2) had significantly prolonged lesion lengths, indicating a faster pathogen spread within the plant tissues. Supporting the limiting effect on the size of lesions, we found that the lesions were significantly shorter in overexpression strains (*OsML1OE-1* and *OsML1OE-2*), indicating that *OsML1* has a significantly negative effect on Xoo pathogen propagation.

OsML1 may thus represent the first identified, with a structurally conserved LPS recognizing structure, functionally defined as reacting to LPS; i.e an identified functional LPS receptor with downstream signalling activity necessary for the innate immunity activation against invading Gram-negative bacteria, such as Xoo, a breakthrough in plant immunity.

### Analysis of the phosphatidylinositol transfer protein family in rice

OsML1 protein contains the PG-PI_TP domain and is homologous to the IV class of the phosphatidylinositol transfer protein family. This family is widely involved in phospholipid transport, membrane signaling pathway regulation, and stress response in plants. The rice whole genome was searched for members of the PITP family, and a total of 30 members were found, including 2 OsML1 genes [2]. Phylogenetic analysis shows that OsML1 and OsML2 are clustered into independent branches in the PITPs phylogenetic tree [2]. (Group I) which is far from the typical SEC14 class members in the system position, indicating that since the OsML1 subgroup is functionally likely MD-2 protein, the OsML1/MD2 could be the earliest MD2 protein evolved, or it could have evolved convergently. Our analysis (**Fig. 5**) shows that there is a fragment duplication relationship between OsML1 and OsML2. This type of duplication often allows new genes to acquire functional innovations through gene rearrangement, structural mutation, or changes in expression regulation [15].

The present study suggests that OsML1 not only obtained a complete structural lipid recognition domain of MD-2 type. The protein shows a wide and strong expression response under various adverse stress conditions, indicating that it has become an adaptive immune regulator. We thus speculate that OsML1 got this enhanced function after the duplication of the *OsML2* gene, followed by adaptive evolution. It thus constitutes a typical example of gene duplication-functional differentiation followed by the evolution of a new function for one of the gene copies [15].

### OsML1 effects on other proteins and the plant metabolism

OsML1 further participates in plant immunity by regulating lipid metabolism. The importance of lipid metabolism in plant immunity is receiving increasing attention, especially the immune signaling molecules involved in regulating the stability of the cell and its membrane structures.

Existing studies have shown that lipid metabolism pathways not only provide the necessary precursors for immune signaling but also regulate the aggregation of receptors on the membrane and the formation of signaling complexes through membrane lipid remodeling, thereby directly or indirectly affecting the plants’ immune responses [16].

In this study, non-targeted metabolomics analysis revealed that 690 types of metabolites in plants underwent significant changes after the OsML1 gene was knocked out. Among them, a significant increase was found in the amount of metabolites such as Beta-Alanine. Sarcosine and AMP may reflect the enhanced signal for plant immune activation, oxidative stress response, or the initiated compensation mechanism. In addition, plant protective metabolites such as lignans and flavonoids, such as Sesamin and Kuwanon C, were significantly reduced in content in the knockout plants, suggesting that the plant’s defense mechanism may be impaired, thereby weakening resistance to pathogens [23,24].

These significantly differently expressed metabolites are mainly concentrated in key pathways such as amino acid metabolism, nucleotide metabolism, lipid metabolism, and antimicrobial metabolism, indicating that OsML1 may optimize the plant’s immune response by regulating the above pathways.

### Possible future research

Future research could further explore the specific molecular mechanisms of OsML1-mediated lipid metabolism and plant immunity, for example, identifying the interaction network between OsML1 and specific lipid metabolism enzymes or signaling components, and analyzing the specific functions of lipid metabolites in immune signal transduction. Doing so could potentially identify the interaction network between OsML1 and specific lipid metabolism enzymes or signaling components, and analyze the specific functions of lipid metabolites in immune signal transduction.

Advanced technologies such as lipidomics, protein interaction omics, and metabolic flow analysis can further be used to clarify how OsML1 regulates membrane microstructure or immune complex formation through lipid metabolism, thereby precisely regulating the intensity and duration of the immune response. or the formation of immune complexes, thereby revealing the regulation of the intensity and duration of the immune response. This will not only help to deepen our understanding of the complex and precise regulation of lipid metabolism and plant immunity, but also provide an important theoretical basis and technical support for improvements of crop disease resistance through metabolic engineering or molecular breeding strategies.

In a long-term perspective, analyzing the relationship between lipid metabolism and plant immunity will greatly promote in-depth and expanded research on plant disease resistance. In particular, a clarification of the detailed mechanism by which OsML1 regulates plant immune response through lipid metabolism pathways is expected to reveal new immune signaling pathways and key nodes by which OsML1 regulates plant immune response through lipid metabolism and PAMPs recognition.

## Conclusion

Rice, as one of the most important food crops in the world, and its immune system research is not only crucially important for crop disease resistance, but also of great significance for investigating the innate immune mechanism of plants.

Our study revealed that the MD-2-like protein OsML1 plays multiple roles, recognizing lipopolysaccharide (LPS), regulating immune responses, and lipid metabolism.

Our discovery study of the MD-2 homologous protein OsML1, with similar functions. provides a breakthrough for the analysis of the LPS sensing mechanism in plants. This potential functional analogy suggests that plants also have evolved an MD-2-like structural module that can recognize exogenous lipid molecules like LPS that can initiate innate immune responses.

## References

[1] Liu J, Liu B, Chen S F, et al. A tyrosine phosphorylation cycle regulates fungal activation of a plant receptor Ser/Thr kinase[J]. Cell Host & Microbe, 2018, 23(2): 241–253.

[2] Pei M, Xie X, Peng B, Chen X, Chen Y, Li Y, Wang Z, Lu G. Identification and Expression Analysis of Phosphatidylinositol Transfer Proteins Genes in Rice. Plants 2023, 12, 2122. 10.3390/plants12112122.

[3] Dow J M, Crossman L, Findlay K, et al. Biofilm dispersal in Xanthomonas campestris is controlled by cell-cell signaling and is required for full virulence to plants[J]. Proceedings of the National Academy of Sciences of the United States of America, 2003, 100(19): 10995–11000.

[4] Meyer A, Puhler A, Niehaus K. The lipopolysaccharides of the phytopathogen Xanthomonas campestris pv. campestris induce an oxidative burst reaction in cell cultures of Nicotiana tabacum[J]. Planta, 2001, 213(2): 214–222.

[5] Darsonval A, Darrasse A, Durand K, et al. Adhesion and fitness in the bean phyllosphere and transmission to seeds of Xanthomonas fuscans subsp. fuscans[J]. Molecular Plant-Microbe Interactions, 2009, 22(6): 747–757.

[6] Gottig N, Garavaglia B S, Garofalo C G, et al. A filamentous hemagglutinin-like protein of Xanthomonas axonopodis pv. citri, the phytopathogen responsible for citrus canker, is involved in bacterial virulence[J]. PLoS One, 2009, 4: e4358.

[7] Jones J D, Dangl J L. The plant immune system[J]. Nature, 2006, 444(7117): 323–329.

[8] Dodds P N, Rathjen J P. Plant immunity: Towards an integrated view of plant-pathogen interactions[J]. Nature Reviews Genetics, 2010, 11(8): 539–548.

[9] Madala N E, Molinaro A, Dubery I A. Distinct carbohydrate and lipid-based molecular patterns within lipopolysaccharides from Burkholderia cepacia contribute to defense-associated differential gene expression in Arabidopsis thaliana[J]. Innate Immunity, 2012, 18(1): 140–154.

[10] Naito K, Taguchi F, Suzuki T, et al. Amino acid sequence of bacterial microbe-associated molecular pattern flg22 is required for virulence[J]. Molecular Plant-Microbe Interactions, 2008, 21(9): 1165–1174.

[11] Bethke G, Unthan T, Uhrig J F, et al. Flg22 regulates the release of an ethylene response factor substrate from MAP kinase 6 in Arabidopsis thaliana via ethylene signaling[J]. Proceedings of the National Academy of Sciences of the United States of America, 2009, 106(19): 8067–8072.

[12] Chinchilla D, Bauer Z, Regenass M, et al. The Arabidopsis receptor kinase FLS2 binds flg22 and determines the specificity of flagellin perception[J]. The Plant Cell, 2006, 18(2): 465–476.

[13] Zipfel C, Kunze G, Chinchilla D, et al. Perception of the bacterial PAMP EF-Tu by the receptor EFR restricts Agrobacterium-mediated transformation[J]. Cell, 2006, 125(4): 749–760.

[14] de Jonge R, van Esse H P, Kombrink A, et al. Conserved fungal LysM effector Eep6 prevents chitin-triggered immunity in plants[J]. Science, 2010, 329(5994): 953–955.

[15] Cisneros AF, Dibyashintan S, Landry CR. Evolutionary causes and consequences of gene duplication. Nature Reviews Genetics, 2026, 10.1038/s41576-026-00935-5.

[16] Raeisi M, Yousefi M, Motavalli R, Hassanzadeh A, Seddighi N, Mehri M, Ebrahimi S, Mehdizadeh A, Abbasi K. Unpacking the lipid–immune axis in health and disease. Immunol Cell Biol, 2026, 104: 208–233. 10.1111/imcb.70076.

